# map3C: a computational tool for processing multiomic single-cell Hi-C data

**DOI:** 10.1101/2025.10.10.681728

**Authors:** Joseph Galasso, Ye Wang, Frank Alber, Jason Ernst, Chongyuan Luo

**Author notes:** Correspondence: Jason Ernst, Chongyuan Luo. **Availability and Implementation:** https://github.com/luogenomics/map3C. **Contact:** Jason Ernst, Chongyuan Luo. **Supplementary information:** supplementary_data.docx.

## Abstract

**Summary:** The emergence of multiomic single-cell Hi-C methods, which simultaneously profile chromatin conformation and other modalities such as gene expression or DNA methylation, creates tremendous opportunities for studying the genome’s structure-function relationships. Existing tools for processing multiomic single-cell Hi-C datasets have certain limitations for downstream bioinformatics analysis. We present map3C, a software tool designed to address these limitations. We demonstrate that map3C improves the quality of multiomic single-cell Hi-C data for analysis and its utility for identifying structural variant locations in the genome.

## Introduction

Genome-wide chromatin conformation, which is the 3D arrangement of chromatin in the cell nucleus, is profiled with various chromatin conformation capture (3C) derived assays such as Hi-C (Lieberman-Aiden *et al*. 2009; Servant *et al*. 2015; Open2C *et al*. 2024). Hi-C uses proximity ligation (PL) to generate DNA fragments containing spatially interacting genomic loci, which are read by high-throughput sequencing (HTS), aligned to a reference genome, and then identified as interactions using software tools such as Pairtools (Open2C *et al*. 2024) and others (Servant *et al*. 2015; Durand *et al*. 2016). These interactions, also called “contacts”, reveal 3D genome features, such as chromatin loops, topologically associating domains, and compartments, which help regulate gene expression (Lieberman-Aiden *et al*. 2009; Dixon *et al*. 2012; Rao *et al*. 2014). In addition to revealing 3D genomic features, the information in Hi-C data has also been shown to be informative for identifying structural variants (SVs) (Wang *et al*. 2020; Song *et al*. 2022; Wang, Luan and Yue 2022).

While originally developed as a bulk assay, recent studies have developed single-cell Hi-C (scHi-C) assays to investigate 3D genomic features at the single cell level (Nagano *et al*. 2013, 2017; Ramani *et al*. 2017). Building upon scHi-C, a number of additional assays have been integrated with other genomic modalities to create “multiomic” assays (Tan *et al*. 2018; Lee *et al*. 2019; Preissl, Gaulton and Ren 2023; Qu *et al*. 2023; Chang *et al*. 2024; Wu *et al*. 2024; Zhou *et al*. 2024a). For instance, LiMCA simultaneously profiles transcription and chromatin conformation (Wu *et al*. 2024), while snm3C-seq simultaneously profiles methylation and chromatin conformation (Lee *et al*. 2019). Multiomic scHi-C creates new bioinformatics challenges and opportunities for integrative analysis. While a few tools have been developed for processing multiomic scHi-C data (Tan *et al*. 2018; Lee *et al*. 2019; Liu *et al*. 2021), challenges still remain.

One challenge is that some multiomic scHi-C assays, such as LiMCA and snm3C-seq, use a specific variation of the PL approach that introduces alignment artifacts in HTS reads (Wu *et al*. 2019). Similar artifacts, though generated by a different mechanism, are also present in single cell methylation data, which motivated the development of a computational method, scBS-map, to remove them (Wu *et al*. 2019). However, no multiomic scHi-C contact-calling tool currently incorporates this functionality. A second challenge is that none of the previously reported contact-calling tools for snm3C-seq data, TAURUS-MH (Lee *et al*. 2019) and YAP (Liu *et al*. 2021) provide quality control (QC) metrics for filtering out low-quality alignments. A third challenge is that no multiomic scHi-C tool calls SVs at 1bp genomic resolution. The feasibility of this for bulk Hi-C data was established with the HiNT-TL tool (Wang *et al*. 2020).

To simultaneously address these challenges, we developed map3C. map3C integrates multiomic scHi-C contact-calling via Pairtools algorithms (Open2C *et al*. 2024) with multimapping trimming, QC metric reporting, and reporting of likely SV breakpoints at 1bp genomic resolution with a variant of HiNT-TL’s algorithm (Wang *et al*. 2020). We present examples demonstrating the utility of these features in the context of multiple LiMCA (Wu *et al*. 2024) and snm3C-seq (Lee *et al*. 2019) datasets from a diverse set of mouse and human tissues.

### Implementation

map3C takes as input paired-end reads locally aligned to a reference genome and produces as output chromatin contacts, which are annotated if they are potentially induced by SVs, and alignments with filtered out mapping artifacts (Fig. 1). Each paired-end read consists of a forward (R1) and a reverse (R2) read from the input DNA sequences (Fig. 1A). If these reads are not bisulfite-converted, map3C requires them to be aligned with BWA MEM (Li 2013) or BWA MEM2 (Vasimuddin *et al*. 2019), as these are recommended aligners by Pairtools (Fig. 1B). However, if the reads were generated by assays that use bisulfite conversion to profile methylation, such as snm3C-seq (Lee *et al*. 2019), map3C requires that the reads be aligned by Biscuit (Zhou *et al*. 2024b) or BSBolt (Farrell *et al*. 2021), which are designed to handle bisulfite conversion and are based on BWA MEM (Fig. 1B). map3C filters out low-quality alignments based on their MAPQ score (Fig. 1C).

**Figure 1.**
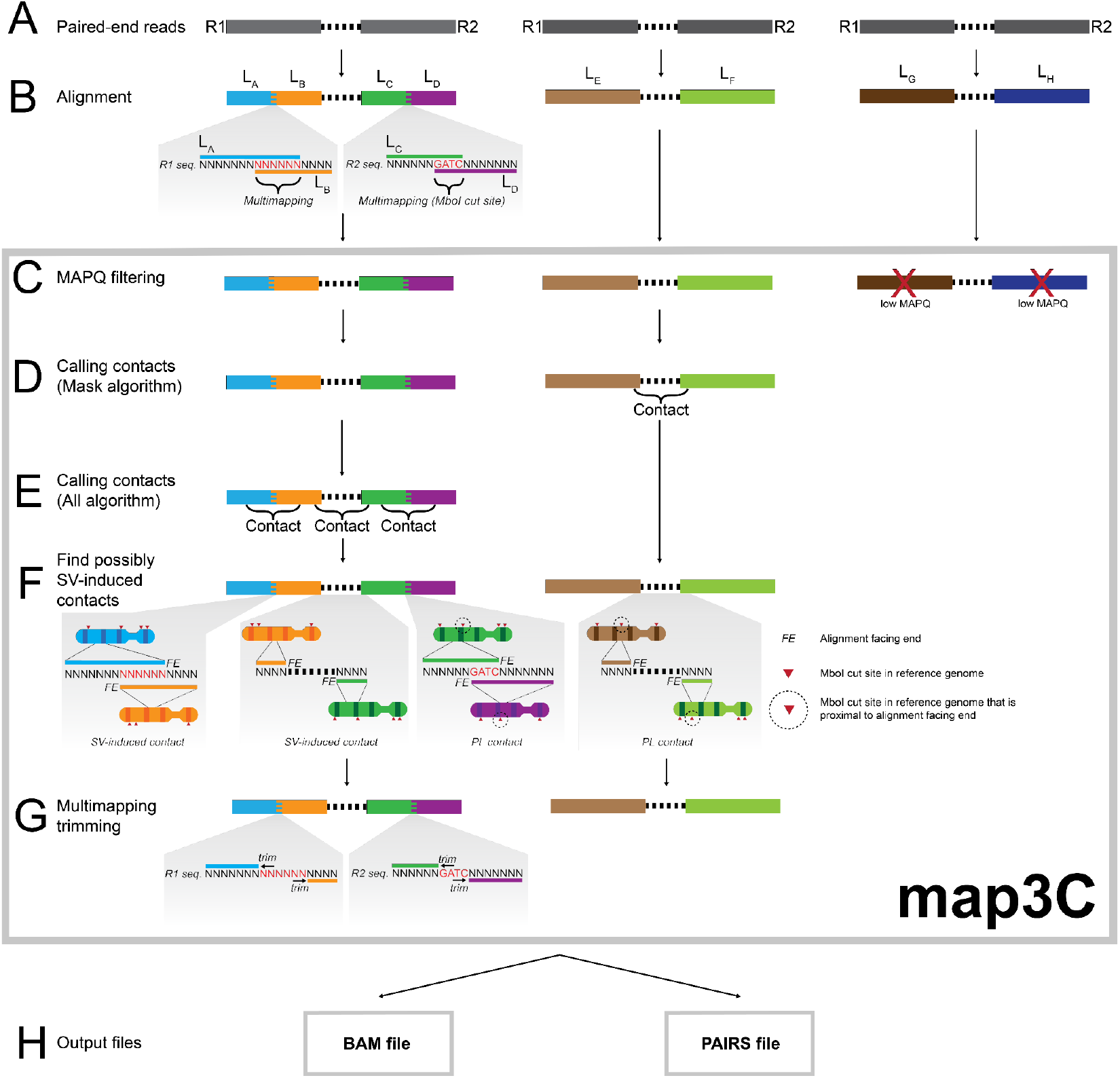
Overview of map3C. A) Paired-end reads for multiomic scHi-C assays. B) Reads are aligned to specific loci (L_A_, L_B_, L_C_, L_D_, L_E_, L_F_, L_G_, L_H_) in the reference genome. For simplicity, these loci are all on different chromosomes. L_A_, L_B_, L_C_, and L_D_ alignments are soft-clipped, and the L_A_/L_B_ and L_C_/L_D_ junctions display multimapping. The L_C_/L_D_ junction’s multimapping is induced by an RE cut site. C) map3C filters out alignments with low MAPQ scores. D) map3C calls contacts with the Pairtools mask algorithm. E) map3C optionally calls contacts with the Pairtools all algorithm. F) map3C analyzes contacts to determine if the alignments’ facing ends are proximal to RE cut sites (represented as red triangles, which are circled if proximal to a facing end). If not, map3C labels these contacts as being more likely induced by SVs. G) Multimapping portions of adjacent soft-clipped alignments are trimmed. H) map3C outputs trimmed alignments in a BAM file and contacts in a PAIRS file, with annotations about RE cut site proximity.

map3C processes alignments through the Pairtools “mask” algorithm (Fig. 1D) and optionally the Pairtools “all” algorithm to generate contacts (Open2C *et al*. 2024) (Fig. 1E). The mask algorithm handles cases where two spatially-interacting genomic loci are present in a read pair, while the all algorithm handles cases where more than two spatially-interacting genomic loci are present (Open2C *et al*. 2024).

map3C finds and annotates contacts that might be caused by SVs (Fig. 1F). These contacts’ alignments are distal from a restriction enzyme (RE) cut site in the reference genome. SVs cause enrichment of contacts distal from RE cut sites, as some read pairs intersect SV breakpoints. Furthermore, if an SV-induced contact’s alignments are adjacent and “soft-clipped”, the junction between the two alignments can reveal the SV breakpoint at 1bp resolution (Rausch *et al*. 2012; Chen *et al*. 2016; Wang *et al*. 2020). Soft-clipped alignments are alignments to only a portion of a read, which are produced by aligners for which map3C is compatible. This approach for identifying contacts that are more likely to be caused by SVs is similar to that of HiNT-TL, but has not been previously integrated into multiomic scHi-C software. Furthermore, map3C reports all contacts that are more likely to be caused by SVs, in contrast to HiNT-TL, which only reports interchromosomal SVs (Wang *et al*. 2020).

map3C removes alignment artifacts at ligation junctions between spatially-interacting genomic loci (Fig. 1G). Some of these artifacts are caused by sticky-end PL, which is used in some multiomic scHi-C assays (Tan *et al*. 2018; Lee *et al*. 2019; Liu *et al*. 2023; Qu *et al*. 2023; Wu *et al*. 2024) since it has efficiency advantages over the alternative blunt-end PL (Lieberman-Aiden *et al*. 2009; Nagano *et al*. 2013, 2017; Rao *et al*. 2014). Sticky-end ligation creates a single RE motif at the junction between two ligated loci. Since each locus borders a motif in the reference genome, aligners may multimap the motif to each locus (Fig. 1A), but the motif can only derive from one locus *in vivo*. map3C removes all multimapping between adjacent soft-clipped alignments from the same read (Fig. 1G) to eliminate inaccurate alignments.

map3C produces two main output files (Fig. 1H). The first is a BAM file containing multimapping-trimmed alignments that are above a minimum MAPQ threshold, which can be customized by the user. The second is a PAIRS file containing contacts (Open2C *et al*. 2024). The contacts in the PAIRS file are annotated if they fit criteria for being more likely to be induced by SVs.

### Example Cases

#### Case 1: Finding SV breakpoints in LiMCA data with map3C

We applied map3C to identify SV breakpoints in K562 cells, a myelogenous leukemia cell line. Specifically, we used map3C to call contacts on 63 K562 cells profiled by LiMCA (Wu *et al*. 2024) to identify 358,716 possible 1bp-resolution SV breakpoints. We note that many of these called breakpoints may not actually be due to SVs and are instead due to technical or alignment artifacts commonly associated with HTS and Hi-C (Imakaev *et al*. 2012; Dixon *et al*. 2018; Koboldt 2020). Thus, to identify higher-confidence 1bp breakpoints, we applied EagleC (Wang, Luan and Yue 2022), a method for calling SV breakpoints at 5kb resolution from Hi-C contact frequencies, to the entire set of K562 contacts, generating 112 breakpoints. The EagleC breakpoints were intersected with the map3C breakpoints to generate a final list of 61 1bp breakpoints, which were within the boundaries of 44 unique EagleC-identified SVs (Fig. S1). Of these 61 1bp breakpoints, 23 were validated by SV calling from whole genome sequencing (WGS) of K562 (Dixon *et al*. 2018). One possible cause of non-validated breakpoints is clonal evolution of K562, which has been reported for other cell lines (Ben-David *et al*. 2018; Petljak *et al*. 2019). A hallmark of breakpoints resulting from sequencing or analysis artifacts is their co-occurrence in multiple LiMCA datasets for different cell lines (Dixon *et al*. 2018; Koboldt 2020). Thus, we determined if the 1bp K562 breakpoints also had evidence of being breakpoints in GM12878, a non-cancer cell line, by applying map3C to LiMCA data for 221 GM12878 cells (Wu *et al*. 2024) (Fig. S1). Of the K562 breakpoints jointly identified with map3C and EagleC, 93% have no supporting GM12878 contacts, despite the GM12878 dataset having 3.1X more HTS reads than the K562 dataset, thus suggesting most breakpoints are not driven by technical artifacts that are common across cell lines.

#### Case 2: Improving alignment accuracy with map3C multimapping trimming

We next demonstrate that map3C multimapping trimming can remove inaccurate alignments. In a LiMCA-profiled mouse olfactory epithelium (MOE) cell (Wu *et al*. 2024), 26% (of *n*=3,342,056) soft-clipped alignments had multimapping, and map3C trimmed out 8% (std=4%) of their spans along their respective reads, on average. Similarly, results were seen in an snm3C-seq profiled mouse embryonic stem cell (mESC) (Lee *et al*. 2019), where 30% (of *n*=251,356) soft-clipped alignments had multimapping and map3C trimmed out 11% (std=7%) of their read spans, on average. To provide evidence that multimapping contributes to inaccurate alignments, we identified LiMCA alignments with a relatively high SNP density and conflicting SNP haplotype assignments (Methods). A potential explanation for the conflicting SNP haplotype assignments in such cases is that the SNP observations from multimapping regions are inaccurate. Consistent with this explanation in two LiMCA-profiled MOE cells (Wu *et al*. 2024), we found 313 alignments that fit the above criteria before map3C multimapping trimming. After trimming, 83% of these conflicts were resolved, as remaining SNPs agreed on a haplotype. For comparison, for each of these alignments, if we assign haplotypes after randomly excluding the same number of SNP observations that were removed by multimapping trimming, the conflict reduction rate drops to 29% on average (std=2%) (Fig. S2).

#### Case 3: map3C QC of snm3C-seq data to enable single-cell Integrative Genome Modeling (IGM)

We show that map3C improves computational 3D chromatin by removing spurious alignments from bisulfite-converted multiomic scHi-C data via MAPQ filtering. To assess contact quality in snm3C-seq, we analyzed the intrachromosomal-to-interchromosomal contact ratio in mESCs. map3C yielded a higher ratio (mean=3.16±1.23) than for TAURUS-MH (mean=0.98±0.26) or YAP (mean=1.10±0.31) on the same dataset (Fig. S3A), closer to the 4.22 ratio reported by the 4DNucleome pipeline (Reiff *et al*. 2022). Improvements from map3C processing are evident in 3D genome structures generated with our modified IGM software (Boninsegna *et al*. 2022). Constructing 3D models that satisfy all chromatin contacts indicates self-consistent, high-quality snm3C-seq data, whereas physically incompatible spurious contacts that cannot be simultaneously realized in 3D increase the fraction of violated constraints per model (restraint-violation rate) and signal poor data quality. Models from map3C-derived contacts showed the lowest restraint-violation rate (Median: 0.004, Mean: 0.006 ±SD: 0.01), outperforming TAURUS-MH (Median: 0.051, Mean: 0.057 ±SD: 0.041) and YAP (Median: 0.037, Mean: 0.039 ±SD: 0.031) (Fig. S3B). TAURUS-MH and YAP-derived structures were excessively compact compared to map3C (Fig. S3C).

With 3D models in hand, we benchmarked structural accuracy against DNA seqFISH+ (Takei *et al*. 2025) imaging data in mESC by comparing (1) the average nuclear radial positioning of genomic regions and (2) interchromosomal contact probability (ICP), a feature linked to transcriptional activity (Boninsegna *et al*. 2022; Yildirim *et al*. 2022, 2023). Models from map3C contacts more closely recapitulated the observed radial profiles (Fig. S3D) and showed substantially better agreement with imaging-derived ICP values compared to YAP and TAURUS-MH (Fig. S3E; Spearman r: map3C = 0.76, YAP = 0.45, TAURUS-MH = 0.51), demonstrating that map3C’s stringent alignment filtering enhances single-cell model fidelity.

## Acknowledgements

We thank Nathan Zemke, Bing Yang, and Matthew Heffel for testing map3C and providing useful feedback. We thank Francesco Musella for discussions and assistance with DNA SeqFISH+ data analysis. We thank all members of the Luo, Ernst, and Alber labs for helpful discussions related to preparation of the manuscript.

## Data Availability

The data underlying this article are available in https://www.github.com/luogenomics/map3C and in its online supplementary material.

