## Supplementary Methods and Figures for "map3C: a computational tool for processing multiomic single-cell Hi-C data"

Joseph Galasso[1,2,3], Ye Wang[1,4,5,6], Frank Alber[4,6], Jason Ernst[1,2,7,8], Chongyuan Luo[1,3]
[1] *Bioinformatics Interdepartmental Program, University of California, Los Angeles, Los Angeles CA 90095, USA*
[2] *Department of Biological Chemistry, David Geffen School of Medicine, University of California, Los Angeles, Los Angeles CA 90095, USA*
[3] *Department of Human Genetics, University of California, Los Angeles, Los Angeles CA 90095, USA*
[4] *Institute of Quantitative and Computational Biosciences (QCBio), University of California Los Angeles, Los Angeles CA 90095, USA*
[5] *Department of Pathology, David Geffen School of Medicine, University of California Los Angeles, Los Angeles CA 90095, USA*
[6] *Department of Microbiology, Immunology, and Molecular Genetics, University of California Los Angeles, Los Angeles CA 90095, USA*
[7] *Department of Computational Medicine, David Geffen School of Medicine, University of California, Los Angeles, Los Angeles CA 90095, USA*
[8] *Computer Science Department, Henry Samueli School of Engineering and Applied Science, University of California, Los Angeles, Los Angeles CA 90095, USA*

### Methods

#### Data sources and preprocessing

For analysis of LiMCA data, FASTQ files for 63 K562 cells, 221 GM12878 cells, and 2 MOE cells were downloaded from NCBI GEO accession GSE240128 and trimmed using the author's reported methods (Wu *et al.* 2024). The MOE cells were from F1 hybrid DBA/2J (JAX 000671) × C57BL/5J mice. For analysis of snmC-seq2 data, 100 mouse brain cells' FASTQ files were obtained from NCBI GEO accession GSE245367 and preprocessed using the YAP trimming approach with Cutadapt v4.9 (GitHub - lhqing/cemba_data: Mapping pipeline for snmC-seq based technologies; Martin 2011; Liu *et al.* 2021; Flint *et al.* 2023). For analysis of snm3C-seq data, 351 mouse embryonic stem cells' (mESC) FASTQ files were obtained from NCBI GEO accession GSE124391 and trimmed using the YAP trimming approach with Cutadapt v4.9 (GitHub - lhqing/cemba_data: Mapping pipeline for snmC-seq based technologies; Martin 2011; Lee *et al.* 2019; Liu *et al.* 2021).

#### Aligning reads

map3C is compatible with alignments of non-bisulfite-converted reads from BWA MEM (Li 2013) and BWA MEM2 (Vasimuddin *et al.* 2019). It is compatible with alignments of bisulfite-converted reads from Biscuit (Zhou *et al.* 2024b) and BSBolt (Farrell *et al.* 2021). For this study, we chose to align non-bisulfite-converted reads generated by LiMCA (Wu *et al.* 2024) with BWA MEM (Li 2013) (Fig. 1B). We chose to align bisulfite-converted reads generated by snm3C-seq and snmC-seq2 with Biscuit (Zhou *et al.* 2024b) (Fig. 1B), as it provides a better alignment strategy for these two assays than BSBolt. This is because these assays have a specific strandedness for bisulfite conversion of reads that can be accounted for by Biscuit. Since snm3C-seq uses a PBAT (Post-Bisulfite Adaptor Tagging) approach for library preparation (Miura *et al.* 2012), R1 reads are aligned to the reverse complementary sequence of the in-silico C to T converted reference genome, whereas R2 reads are aligned directly to the in-silico converted reference. Since Hi-C read pairs can contain distal genomic loci, all aligners are run with the -P and -S parameters, which lead the aligners to map R1 and R2 independently.

We used the hg38 reference genome for human data and the mm10 reference genome for mouse data, including all ALT contigs. We downloaded these references from the UCSC genome browser. For aligning snm3C-seq and snmC-seq2 data, we added the lambda phage DNA genome sequence as a contig into the reference genome, as this phage is commonly used to ascertain bisulfite-conversion efficiency in bisulfite-converted libraries, although the libraries we used did not include it. The lambda phage genome was downloaded from GenBank (https://www.ncbi.nlm.nih.gov/nuccore/J02459.1?report=fasta).

map3C retains all alignments that are above a minimum mapping quality (MAPQ) threshold, which we set to 30 (Fig. 1C). This is the same threshold as the default threshold used by Juicer (Durand *et al.* 2016) for higher quality Hi-C contacts. Further justification for this threshold comes from analyzing alignments from snmC-seq2, which performs whole-genome bisulfite sequencing (WGBS) with no Hi-C component. As there is no Hi-C component, read pairs are expected to only contain alignments to the same chromosome (cis pairs) and thus read pairs with alignments to different chromosomes (trans pairs) are more likely to be mapping artifacts. We observed that a minimum MAPQ of 30 effectively balances reducing trans pairs and increasing cis pairs (Fig. S4A&B).

After this MAPQ filtering, we observed that Biscuit sometimes multimaps a portion of a given read with at least two soft-clipped alignments, which are alignments of less than the full span of the original read to a genomic locus. This multimapping phenomenon is rare, affecting only a small fraction of reads (mean=0.02%, standard deviation=0.003%) in snm3C-seq-profiled mouse embryonic stem cells (mESC). map3C addresses this issue of multimapping by only retaining the alignment with the highest MAPQ score for a given portion of a read that is covered by multiple alignments. If there is a tie for highest MAPQ score, then map3C retains the alignment with the most mapped nucleotides. If there is still a tie for length, then map3C chooses an alignment randomly.

**Calling contacts with map3C**

map3C passes alignments into the Pairtools “mask” (Fig. 1D) and “all” (Fig. 1E) algorithms (Open2C *et al.* 2024b) to call contacts, where contacts are pairs of alignments from the same read pair. The Pairtools mask algorithm is attempted first, which is capable of calling contacts on read pairs with a single ligation (Fig. 1D). Any read pair that produces no contacts from the Pairtools mask algorithm is then optionally processed by the Pairtools all algorithm (Fig. 1E), which identifies read pairs that contain multiple ligation events. We used this option on all data processed in this study to be inclusive of multi-way contacts. We modified the Pairtools all algorithm to optionally report contacts for all possible combinations of alignments in a read pair, thereby including alignment pairs that were not directly ligated to each other. This can be useful for finding multi-way chromatin interactions, which is of interest to some single-cell Hi-C studies, such as GAGE-seq (Zhou *et al.* 2024a), although we did not consider these interactions in our analysis.

**Annotating contacts with map3C**

map3C annotates contacts if their two alignments’ genomic loci are close to RE cut sites in the reference genome (Fig. 1F), indicating that they are more likely to have been generated by PL. It should be noted that since PL joins loci that are distal in the 1D genome, each alignment can be proximal to a different cut site. The criteria for making the determination of whether a contact is close to a RE cut site differs if the contact involves alignments from different (R1 or R2) reads or the same read. In the former case, we use the method from (Jin *et al.* 2013), which requires a cut site to be within 500bp of the facing side of each alignment, which is roughly half of a 1000 bp sequencing insert. In this case, map3C only looks for cut sites in the appropriate genomic direction, given the strandedness of the alignments. For example, if two proximal soft-clipped alignments both align to the watson (+) strand, the R1 alignment’s 3’ side must have a downstream cut site within 500bp. On the other hand, the R2 alignment's 5' side must have an upstream cut site within 500bp. This parameter can be modified depending on the sequencing insert size distribution.

If a contact involves two adjacent soft-clipped alignments from the same read, map3C uses a modified procedure to the approach used in the case that the alignments are from different reads. The first modification is the nearest RE cut site is chosen for each alignment without consideration of direction. The reason for this modification is the possible presence of multimapping at the junction of the alignments. This multimapping is defined formally as follows. To describe the location of a bp in the read, the first bp at the 5' end of a read is denoted as $r_5 = 1$. The bp at the 3' end of the read is denoted as the length of the read, $r_3$. The start and end of a soft-clipped alignment's span on the read occurs at positions $r_s$ and $r_e$, where $r_5 \leq r_s < r_e \leq r_3$. This soft-clipped alignment may have adjacent soft-clipped alignments. An upstream soft-clipped alignment's span is $[r_{5,s}\ , r_{5,e}]$, whereas a downstream alignment's span is $[r_{3,s}\ , r_{3,e}]$. Multimapping occurs when $r_{5,e} \geq\ r_s$ or $r_{3,s} \leq\ r_e$. In the former case, the span of the multimapped region of the read would be $[r_s\ , r_{5,e}]$ and in the latter case, the span of this region would be $[r_{3,s}\ , r_e]$. With regards to assigning RE cut sites to adjacent soft-clipped alignments, this multimapping could cause the nearest cut site to be either upstream or downstream of the facing side of an alignment locus.

The second modification in the case of two adjacent soft-clipped alignments from the same read is the use of a shorter maximum distance threshold. We empirically set this threshold to 20bp by examining the distance distribution from soft-clipped alignments to the nearest RE cut site in snm3C-seq vs snmC-seq2 data. We found that snm3C-seq alignments were substantially concentrated up to approximately 20 bp, whereas snmC-seq2 alignments had a relatively uniform distribution across distances well past 20bp (Fig. S5). Conveniently, this also matches the Pairtools default gap size parameter of 20bp (Open2C *et al.* 2024b), ensuring that RE cut sites are not separated from alignments by large gaps. If one assumes that all alignments accurately represent the *in vivo* locations HTS reads observe, this parameter should be 0bp, as soft-clipped alignments at a PL site should directly border a cut site. However, the presence of gaps and multimapping between adjacent soft-clipped alignments motivates an increased threshold. Consequently, multiple cut sites may be proximal to an alignment. To handle this issue, the cut site that is the closest to the alignment is assigned to it. If there is a tie between two cut sites, one is randomly selected.

**Multimapping and RE cut site trimming**

map3C trims adjacent soft-clipped alignments to remove multimapping between them (Fig. 1G). To eliminate this multimapping, we use an approach similar to scBS-map (Wu *et al.* 2019), where the multimapping portion of each soft-clipped alignment is trimmed, generating new spans for each alignment, such that $r'_{5,e} < r'_s < r'_e < r'_{3,s}$, using the above terminology. Specifically, if $r_{5,e} \geq\ r_s$ originally, then map3C sets $r'_{5,e} =\ r_s - 1$ and $r'_s =\ r_{5,e} + 1$. If $r_{3,s} \leq\ r_e$ originally, then map3C sets $r'_{3,s} =\ r_e + 1$ and $r'_e =\ r_{3,s} - 1$. Unlike scBS-map, which trims 10bp by default off of each soft-clipped alignment's ends, map3C's approach computes and removes the specific number of multimapped bp for each soft-clipped alignment (Wu *et al.* 2019).

While the above algorithm fully removes multimapping of each alignment, there are two particular cases where it does not create accurate boundaries between the alignments. In these cases, the adjacent soft-clipped alignments' facing ends are each proximal to an RE cut site in the reference genome, suggesting that their loci were ligated by PL. The first case occurs when the flanking sequence around the RE cut site motif may be highly similar in each ligated locus. Thus, the multimapping portion of each alignment could be longer than the RE cut site motif. After application of the multimapping trimming algorithm, the facing ends of alignments might not directly border a RE cut site, which is the true *in vivo* boundary between the loci (Fig. S6A). The second case occurs if no multimapping is present between the soft-clipped alignments, but either of the alignments intersects with its respective RE cut site motif. Thus, the multimapping trimming algorithm would not be applied to these alignments, but at least one's facing end would extend over its proximal RE cut site (Fig. S6B).

map3C detects these two cases using the aforementioned RE cut site assignments to alignments and addresses these cases with an alternative trimming algorithm (Fig. S6). Using the previous notation, a soft-clipped alignment has a span of $[r_s, r_e]$ and an upstream soft-clipped alignment has a span of $[r_{5,s}, r_{5,e}]$. Let the upstream alignment's proximal RE cut site to its 3' end span $[r_{5,a}, r_{5,b}]$, where $r_{5,a} < r_{5,b}$, and let the first alignment's proximal RE cut site to its 5' end span $[r_a, r_b]$, where $r_a < r_b$. If $r_b \geq r_s$, then $r'_s = r_b + 1$, and if $r_{5,a} \leq r_{5,e}$, then $r'_{5,e} = r_{5,a} - 1$. For first of these two cases, there is a possibility that multimapping remains if RE cut site assignments or alignments are not accurate; however, we found that this was rare, occurring in a small fraction (M=0.98%, SD=0.010%) of contiguous soft-clipped alignment pairs in mESC cells' mESC data. If this happens to be the case, then the default multimapping trimming algorithm is subsequently applied to fully eliminate residual multimapping.

**Removing proximal intrachromosomal contacts**

map3C filters all contacts to remove "dangling end" artifacts from Hi-C assays (Belton *et al.* 2012), which are caused by sequencing inserts without ligations. These are especially prevalent in many single-cell Hi-C assays (Tan *et al.* 2018; Lee *et al.* 2019; Liu *et al.* 2023; Wu *et al.* 2024; Zhou *et al.* 2024a), which, unlike traditional Hi-C protocols, do not use streptavidin beads for enrichment of biotin-labeled DNA fragments created by PL. To remove dangling end artifacts, map3C requires by default that contact loci must either be 1) interchromosomal or 2) intrachromosomal and at least 1000 bp apart. This distance was selected given the previously discussed expectation for sequencing inserts (Jin *et al.* 2013). TAURUS-MH utilized this threshold (Lee *et al.* 2019), whereas YAP used 2500 bp (GitHub - lhqing/cemba_data: Mapping pipeline for snmC-seq based technologies).

**Output files of map3C**

map3C writes output contacts in the “pairs” (Lee *et al.* 2022) format, like Pairtools (Open2C *et al.* 2024b), where each row contains information for one contact (Fig. 1H). In the third and fifth columns of a pairs file, which are set to report a single genomic position for the first and second alignment in a contact, respectively, map3C reports the position of each alignment’s side that faces the other alignment in the contact. This enables map3C to report the precise breakpoints for SVs. map3C takes advantage of the flexibility of the pairs format to add extra columns after set columns, which note if a given contact has 1) assigned restriction enzyme cut sites and 2) if the contact was formed by contiguous soft-clipped alignments. Finally, map3C generates a BAM file that contains only alignments that meet the MAPQ threshold and do not contain multimapping (Fig. 1H).

**Processing of multiomic scHi-C and snmC-seq2 data with map3C**

map3C was used to generate contacts for the snmC-seq2 mouse brain cells (Flint *et al.* 2023); snm3C-seq (Lee *et al.* 2019) mESC cells; and LiMCA (Wu *et al.* 2024) MOE, K562, and GM12878 cells. TAURUS-MH and YAP were run on the snm3C-seq data with default parameters. We removed PCR duplicates from map3C-generated contacts with Pairtools dedup (Open2C *et al.* 2024b). Pairtools dedup was not applied to TAURUS-MH- and YAP-generated contacts, as these pipelines have internal duplicate removal algorithms. Pairtools highcov was applied to remove contacts from genomic regions with higher coverage than would be expected given the diploid copy number of single cells (Open2C *et al.* 2024b). As mentioned before, snmC-seq2 “contacts” are not caused by PL and are more likely to be the result of mapping artifacts, but are useful for benchmarking map3C performance and previously discussed analyses of important map3C parameters.

**SV calling**

K562 and GM12878 LiMCA data were processed with map3C. PCR duplicates and high coverage contacts were filtered out with Pairtools. Pseudobulk contact matrices for all K562 cells and GM12878 cells were separately generated with Cooltools (Open2C *et al.* 2024a). K562 SVs were identified with EagleC (Wang, Luan and Yue 2022; Wu *et al.* 2024). K562 and GM12878 contacts caused by soft-clipped alignments and not proximal to RE cut sites as determined by map3C were aggregated separately, creating a set for K562 and a set for GM12878. The K562 set was intersected with EagleC SV boundaries. Specifically, for a contact to intersect with an SV, each locus of the contact had to fall within the corresponding 5kb loci reported by EagleC. The contact loci were reported as a possible bp-resolution breakpoint for the SV. As multiple contacts may indicate the same bp-resolution breakpoint, the sum of contacts supporting each unique bp-resolution breakpoint is reported (Fig. S1). Each K562 bp-resolution breakpoint was intersected with the GM12878 set to determine if the breakpoint was also seen in GM12878. A GM12878 contact was reported to overlap a K562 breakpoint if the contact’s two associated genomic loci were each within ±10bp from the corresponding K562 breakpoint locus. The same criteria was used to intersect the K562 bp-resolution breakpoints with K562 SVs identified with WGS (Dixon *et al.* 2018).

**Assessing map3C multimapping trimming performance with haplotype phasing**

For computing how much of alignments are trimmed by map3C in LiMCA and snm3C-seq data, we defined each alignment's span on the read as before, with $[r_s, r_e]$. After trimming, the new span is $[r'_s, r'_e]$. The percent of the original alignment span that is removed is computed as $\frac{(r_e - r'_e) + (r'_s - r_s)}{r_e - r_s + 1}$.

Alignment haplotype phasing of multiomic scHi-C data has been demonstrated previously with the hickit software package (GitHub - lh3/hickit: TAD calling, phase imputation, 3D modeling and more for diploid single-cell Hi-C (Dip-C) and general Hi-C) and an algorithm from (Tan *et al.* 2018). We incorporated this algorithm into map3C to analyze the impacts of multimapping trimming on haplotype phasing. In map3C's implementation, the locus of each alignment is intersected with a database of known phasing of SNPs that is provided as input. If the alignment's SNP observations unanimously agree with one haplotype, it is assigned that haplotype. Otherwise, it is not phased due to conflicting haplotype information. If the HTS alignments are bisulfite-converted, map3C ascertains the alignment's displayed conversion pattern (C>T or G>A) in the BAM file with the alignment's "YD" BAM tag for Biscuit or the alignment's "YS" BAM tag for BSBolt. If a SNP's reference and alternate allele show the expected conversion pattern, then this SNP is not used for phasing, as its alternate allele may have been the result of bisulfite conversion (Wu *et al.* 2024). Prior to applying this algorithm to the LiMCA MOE alignments (Wu *et al.* 2024), we first filtered out alignments to ALT contigs and alignments that intersected with the mm10 excluded regions (https://github.com/Boyle-Lab/Blacklist/raw/refs/heads/master/lists/mm10-blacklist.v2.bed.gz) (Amemiya, Kundaje and Boyle 2019). The MOE phased SNP database was obtained from the LiMCA authors (https://github.com/zhang-jiankun/LiMCA/raw/refs/heads/main/data/mouse_snps/dba2.txt.gz) (Tan *et al.* 2018).

To compare the likelihood of reducing phasing conflicts using multimapping trimming vs. randomly eliminating SNPs from alignments (Fig. S2), we first subset all alignments (MAPQ≥30) with ≥ 3 overlapping SNPs and a conflicting phase assignment prior to trimming. We required ≥ 3 overlapping SNPs because we wanted to reduce the likelihood of instances where trimming removes all but one SNP observation, which will eliminate corroborating information for phasing assignment. For each alignment, we counted the number of SNP observations that would be removed as a consequence of multimapping trimming and determined if the alignment could be phased. We then randomly removed that same number of SNPs and determined if the alignment could be phased. The fraction of all alignments with resolved phasing conflicts through random SNP removal was computed. The randomization process was repeated 1000 times, leading to 1000 conflict resolution fractions, the distribution of which is shown in Fig. S2.

**Comparing map3C, TAURUS-MH, and YAP cis:trans ratios**

To compare cis:trans ratios of contacts between TAURUS-MH, YAP, and map3C (Fig. S3A), we defined cis contacts as those that are intrachromosomal and have a distance >20kb apart. This was to more definitively eliminate dangling ends and focus on contacts that are of more interest in studying long-range chromatin structure (Ramani *et al.* 2017).

**IGM modeling**

For genome representation, chromosomes are segmented into genomic regions of 200-kb DNA sequence length, each represented by chromatin domain with spherical volume, defined by an excluded volume with a radius r = 112nm, which guarantees a 20% volume occupancy of the diploid genome in the nucleus (Tjong *et al.* 2016). The nuclear shape is modeled as a sphere with radius 4,500nm (Milo *et al.* 2010).

In a diploid genome, each autosome genomic region has two homologous copies, and the diploid genome is thus represented by a total of $N = 25965$ chromatin domains. Chromosomes are modeled as polymer chains subject to chain connectivity, chromatin contacts, excluded volume and nuclear volume restraints.

We represent the population of all genome structures in *S* cells as $X = \{X_1, X_{2,} \ldots, X_S\}$, with each single-cell genome structure $X_s = \{\vec{x}_{is} \in R^3, i = 1,2, \ldots, N\}$ as a set of 3D vectors representing the center coordinates of each chromatin domain with $N$ as the total number of chromatin domains in the diploid genome.

The scHi-C data is expressed as a third-order tensor $A = (a_{IJs})_{H\times H\times S}$ where $a_{IJs} \in \{0,1,2\}$ indicates the number of contacts between the genomic regions $I$ and $J$ in cell $s$. Contact counts greater than 2 are capped at 2 to exclude spurious inflation of contact multiplicity caused by fragment-level redundancy within 200 kb bins. The variable $H$ denotes the total number of genomic regions when homologous copies are not distinguished. By convention, capital indices (e.g., $I$,$J$) refer to regions without distinguishing between two homologous copies, while lowercase indices (e.g., $i, i$' and $j, j$') distinguish between them.

An ambiguity of scHi-C data is that, while it records which regions $I$ and $J$ are observed in contact in each cell, it does not specify which homologous chromosome copies realizes these contacts. Thus, contact tensor $A$ is incomplete: it lacks information about whether the observed contact involves homologous domain copies $i$ or $i$' for region $I$, and $j$ or $j$' for region $J$. To resolve this ambiguity in an optimization process we introduce a latent contact indicator tensor $W = (w_{ijs})_{N\times N\times S}$, a binary-valued third-order tensor that specifies contacts between chromatin domains $i$ and $j$ for each homologous copy in each cell. Here, $w_{ijs} = 1$ indicates that a contact between a specific pair of homologous loci $i$ and $j$ is present in structure $s$, while $w_{ijs} = 0$ indicates its absence.

We aim to infer the ensemble of single-cell diploid 3D genome structures consistent with all scHi-C data $A$, by jointly optimizing the population of structures $X$ and the latent contact assignments $W$ that best reconciles the ambiguity in the scHi-C data. Following our IGM framework, this is formulated as a maximum likelihood estimation problem and solved iteratively with a variant of the expectation-maximization algorithm, combined with optimization strategies for scalability and efficiency as described in (Tjong *et al.* 2016; Boninsegna *et al.* 2022).

Each iteration consists of two steps. In the assignment step (A-step), chromatin contacts are optimally allocated in $W$ across all structures ($X$) by maximizing the log-likelihood given the fixed structures from the previous iteration (Boninsegna *et al.* 2022). In the modeling step (M-step), 3D genome structures $X$ are generated by imposing physical contact restraints specified in $W$, while satisfying polymer chain connectivity, excluded volume, and nuclear confinement. This is achieved by using molecular dynamics (MD) simulated annealing (Kirkpatrick, Gelatt and Vecchi 1983) followed by conjugate gradient minimization (Hestenes and Stiefel 1952; Boninsegna *et al.* 2022).

To ensure robust convergence, scHi-C contacts are incorporated gradually in a stepwise process. Contacts are ranked by their pseudo-bulk probability $P(I,J)$, defined as the fraction of cells a contact $(I,J)$ is observed. At stage $t$, only contacts with $P \geq \theta_t$ are included, and the A/M optimization is run until convergence. Convergence is assessed by monitoring normalized residuals (Boninsegna *et al.* 2022): a restraint is considered violated if its residual exceeds 1, and a stage is considered converged when the global violation rate falls below 0.01%. After convergence, the threshold $\theta_t$ is lowered, additional contacts are added, and the A/M cycle is repeated until the final probability threshold $\theta_t$=0.008 is reached and all structures converge. The thresholds used in our modeling protocol were [1, 0.2, 0.1, 0.05, 0.02, 0.01, 0.008].

As shown previously, gradual increase of the optimization hardness in a stepwise fashion can effectively guide the search for the best solution and facilitates the detection of cooperative chromatin interactions (Boninsegna *et al.* 2022; Yildirim *et al.* 2022, 2023).

### Supplementary Figures

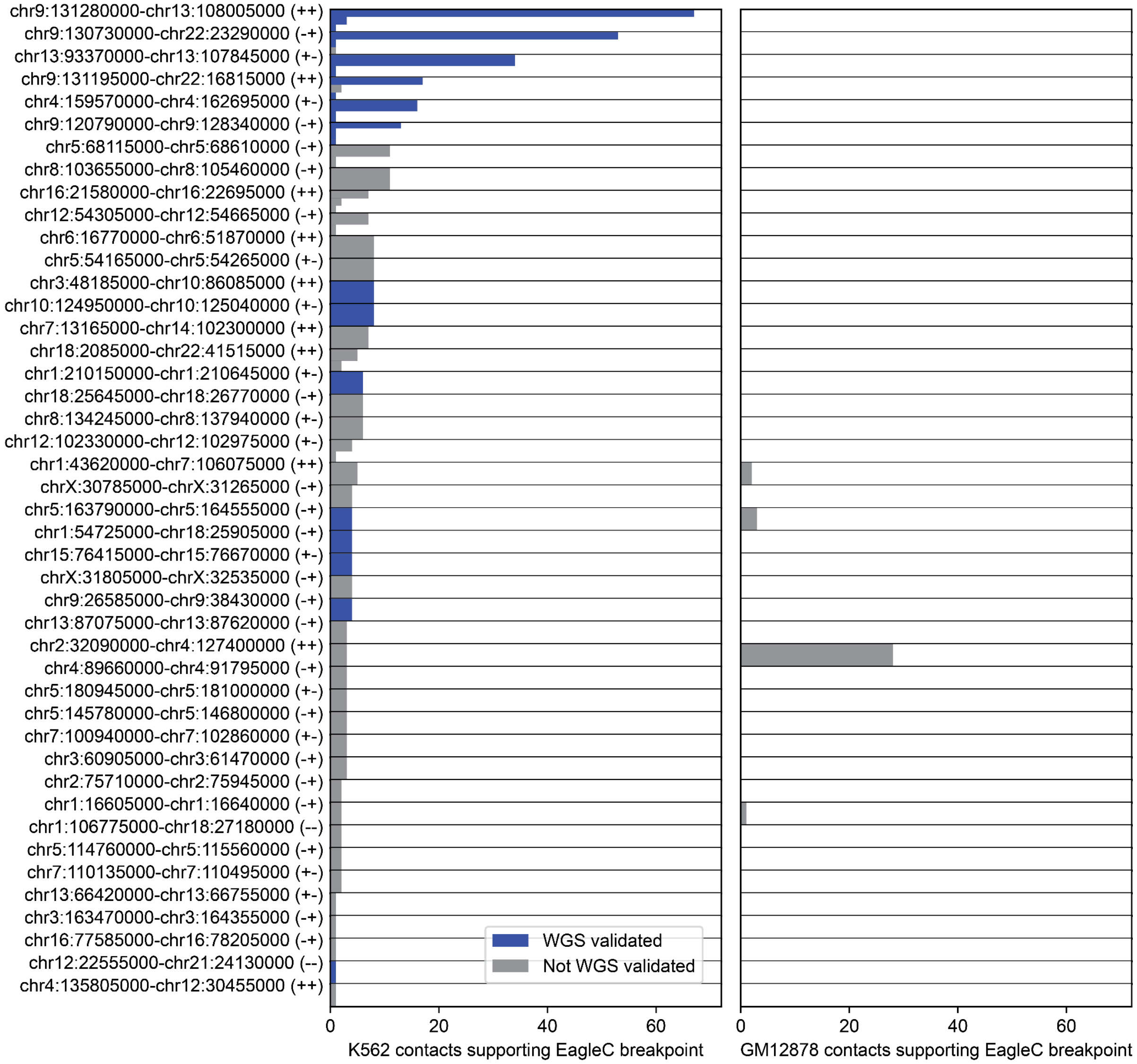

**Supplementary Figure 1. map3C identifies 1bp K562 SV breakpoints.** On the left, EagleC 5kb breakpoints for K562 pseudobulk SVs are reported on the y-axis. These breakpoints must intersect at least one map3C K562 contact that is distal from RE cut sites. For each EagleC breakpoint, the x-axis shows the total number of intersecting contacts that are distal from RE cut sites. The number of contacts is stratified by unique 1bp map3C breakpoints, since EagleC breakpoints can intersect multiple 1bp map3C breakpoints. Each unique 1bp breakpoint is annotated if it is supported by WGS of K562. On the right, the number of map3C GM12878 contacts that are distal from RE cut sites and support each 1bp K562 breakpoint are shown.

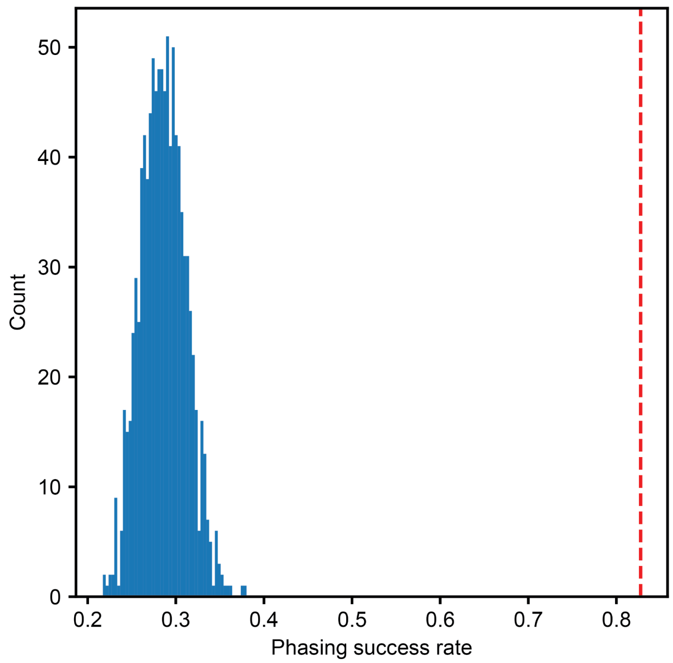

**Supplementary Figure 2. map3C multimapping trimming reduces soft-clipped alignment haplotype phase conflicts.** The red line indicates the fraction of conflicts that are resolved by multimapping trimming. The blue histogram shows the distribution of 1000 conflict-resolution fractions that were generated by randomly removing SNPs.

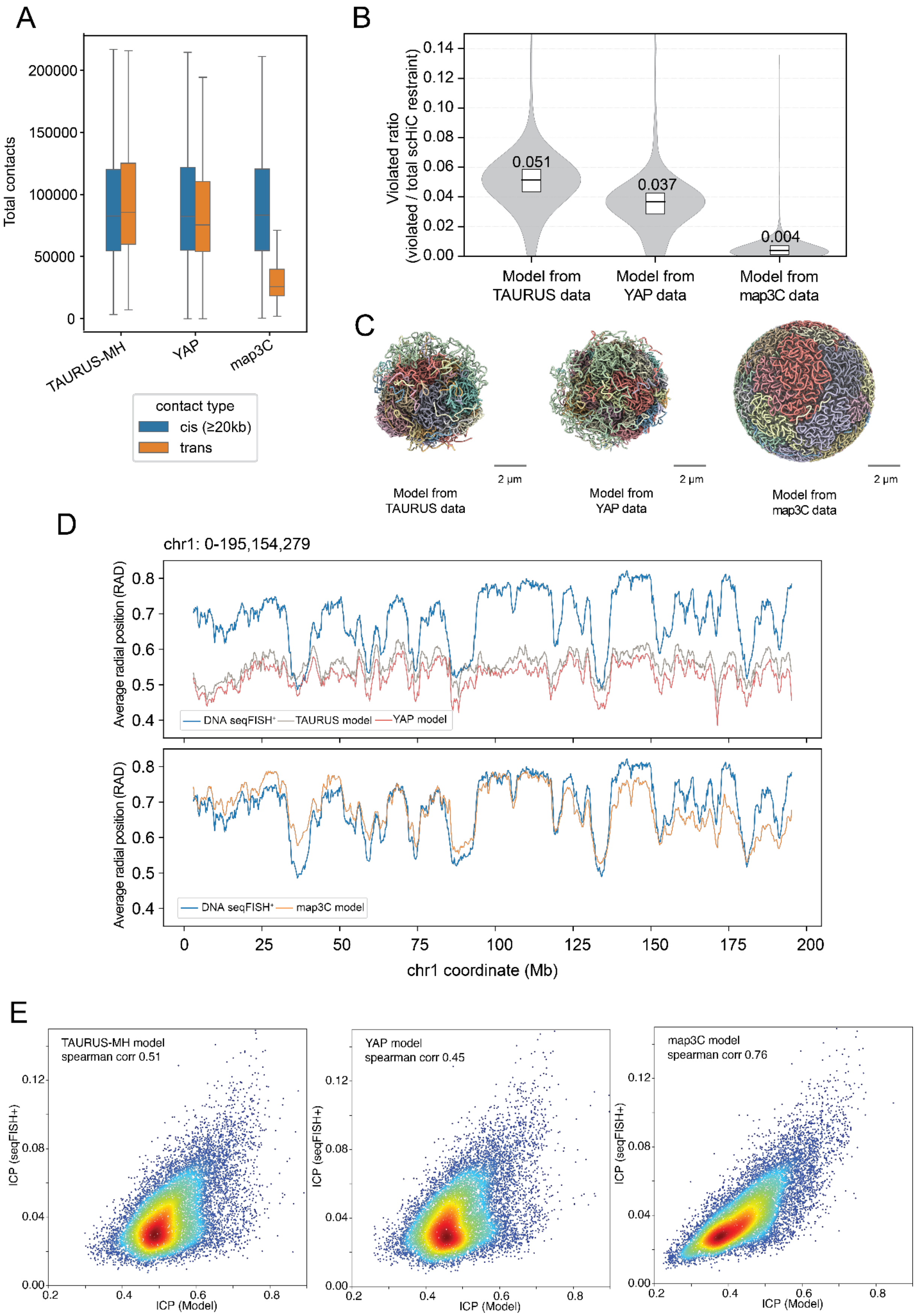
A
Total contacts
200000
150000
100000
50000
0
TAURUS-MH
YAP
map3C
contact type
cis (≥20kb)
trans
B
Violated ratio
(violated / total scHiC restraint)
0.14
0.12
0.10
0.08
0.06
0.04
0.02
0.00
0.051
0.037
0.004
Model from TAURUS data
Model from YAP data
Model from map3C data
C
2 μm
Model from TAURUS data
Model from YAP data
Model from map3C data
D
chr1: 0-195,154,279
Average radial position (RAD)
DNA seqFISH⁺
TAURUS model
YAP model
map3C model
chr1 coordinate (Mb)
E
TAURUS-MH model
spearman corr 0.51
YAP model
spearman corr 0.45
map3C model
spearman corr 0.76
ICP (seqFISH+)
ICP (Model)

**Supplementary Figure 3. map3C QC of snm3C-seq data leads to improved cis:trans ratios and IGM models** A) Comparison of snm3C-seq trans and cis contacts generated by TAURUS-MH, YAP, and map3C across 351 cells. B) Distribution of the violated-restraint fraction (violated/total scHi-C restraints per cell) for IGM chromatin models generated with TAURUS-MH, YAP, and map3C contacts. C) Visualization of representative 3D genome structure models generated with IGM using TAURUS-MH, YAP, and map3C contacts. D) Comparison of average radial position profile of chromosome 1 between DNA seqFISH+ imaging data and genome structure models generated with TAURUS-MH, YAP (upper panel) and map3C contacts (lower panel). Radial positions were averaged across all single cells and normalized on a scale from 0 (nuclear center) to 1 (nuclear envelope). E) Correlation between imaging-derived and model-derived inter-chromosomal contact probability (ICP) for IGM models generated with TAURUS-MH, YAP, and map3C contacts.

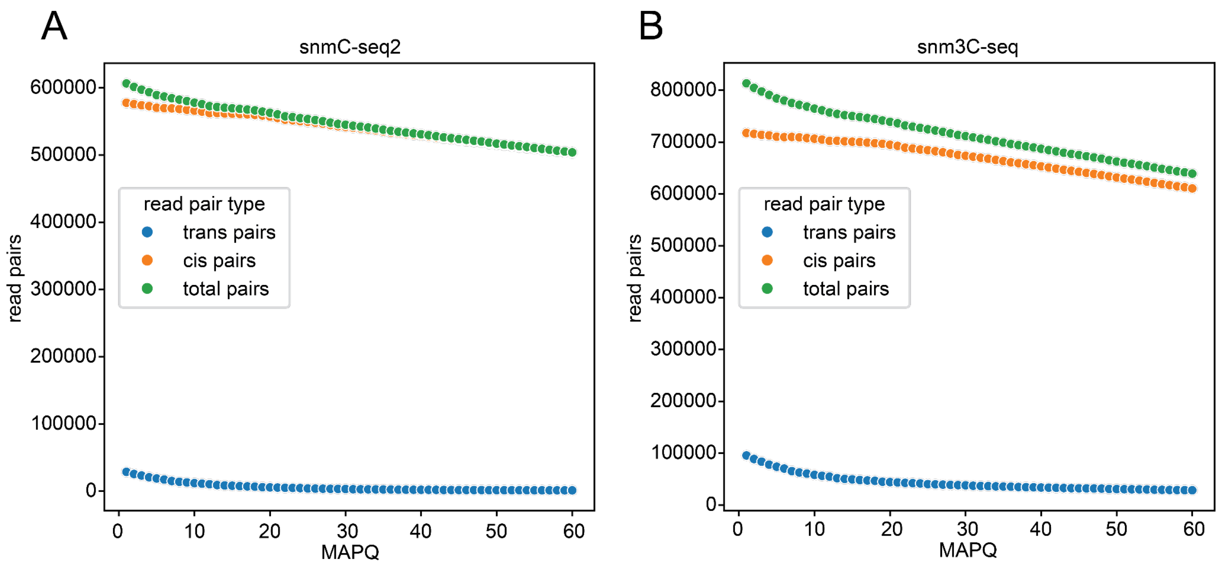

**Supplementary Figure 4. Read pair alignments decrease as MAPQ threshold is increased.** For (A) snmC-seq2 mouse brain data and (B) snm3C-seq mESC data, the number of read pairs with ≥2 alignments that exclusively align to one chromosome (cis), the number of read pairs with ≥2 alignments that align to multiple chromosomes (trans), and the sum of these values are contrasted as MAPQ increases.

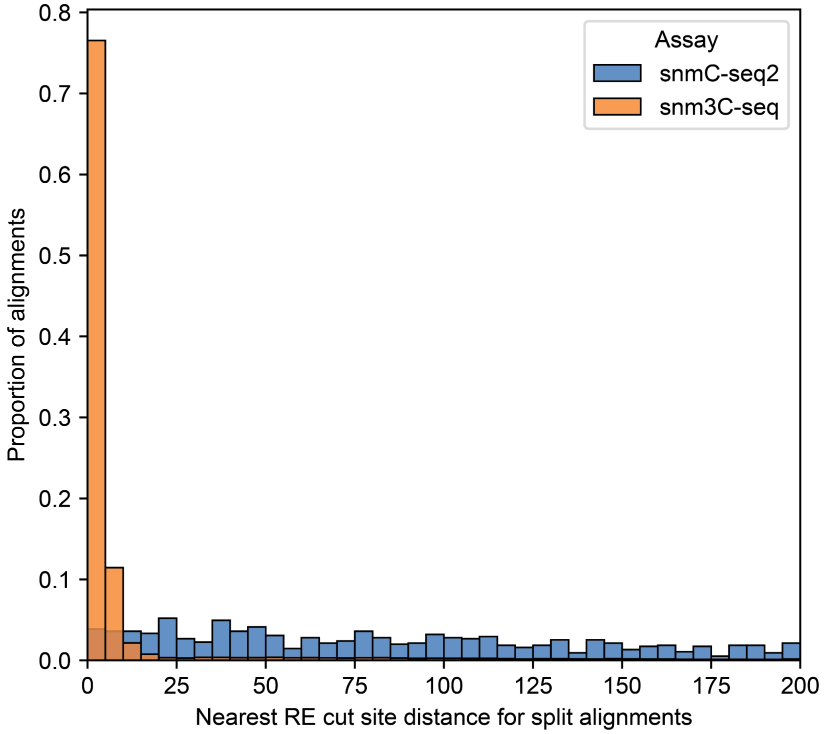

**Supplementary Figure 5. snm3C-seq soft-clipped alignments show proximity to RE cut sites relative to snmC-seq2 soft-clipped alignments.** Distribution of distance from soft-clipped alignments to the nearest RE cut site in one snmC-seq2 mouse brain cell (n=1,230 alignments) and one snm3C-seq mESC cell (n=49,696 alignments). These alignments all face an adjacent soft-clipped alignment from the same read, and the distance is measured from the end of the alignment that faces the adjacent alignment.

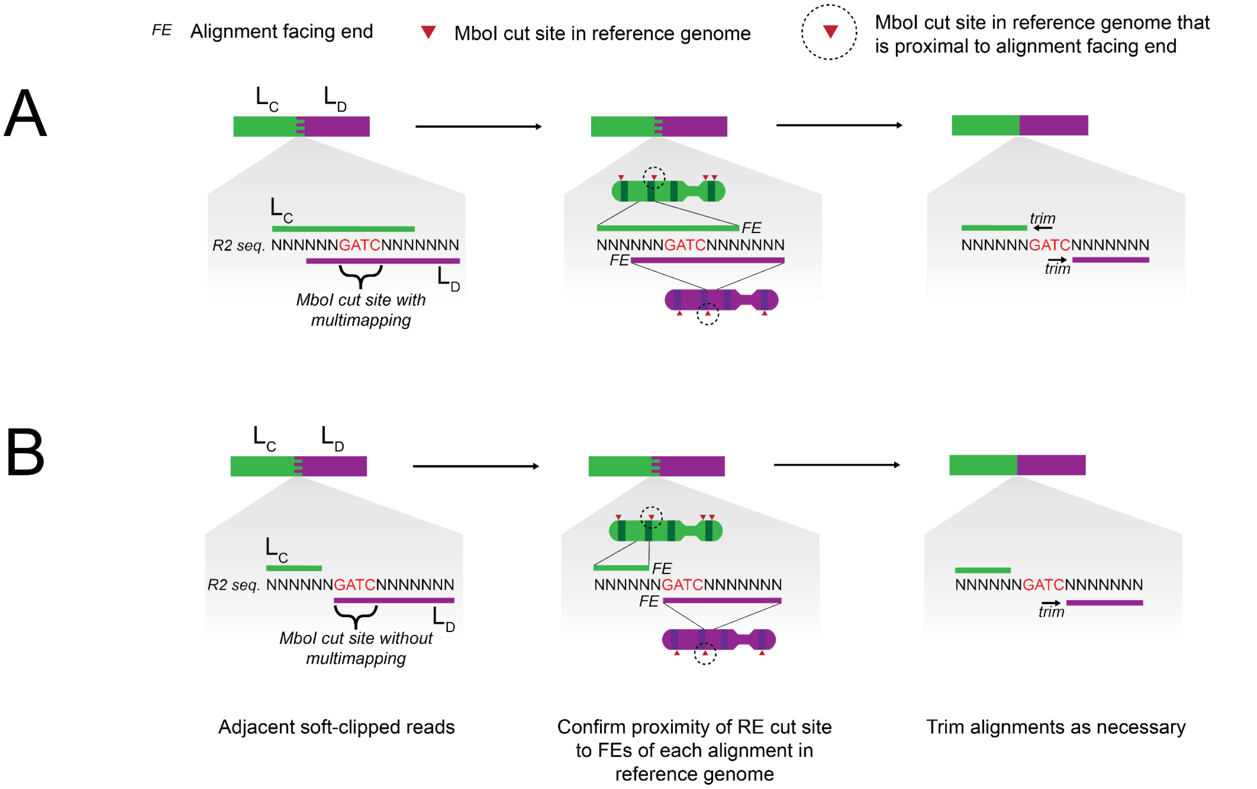

**Supplementary Figure 6. Trimming alignments proximal to RE cut sites.** A) In the case that multimapping is present between alignments, they are each trimmed such that they fully exclude their respective proximal RE cut site loci in the reference genome. B) In the case that multimapping is not present between alignments, it is possible that one or more still includes its respective proximal RE cut site locus in the reference genome. Alignments that fit this criteria are trimmed to fully exclude their RE cut sites.

GitHub - lh3/hickit: TAD calling, phase imputation, 3D modeling and more for diploid single-cell Hi-C (Dip-C) and general Hi-C. *GitHub*.

GitHub - lhqing/cemba_data: Mapping pipeline for snmC-seq based technologies. *GitHub*.

Hestenes MR, Stiefel E. Methods of conjugate gradients for solving linear systems. *J Res Natl Bur Stand (1934)* 1952;**49**:409.

Jin F, Li Y, Dixon JR *et al.* A high-resolution map of the three-dimensional chromatin interactome in human cells. *Nature* 2013;**503**:290–4.

Kirkpatrick S, Gelatt CD Jr, Vecchi MP. Optimization by simulated annealing. *Science* 1983;**220**:671–80.

Lee D-S, Luo C, Zhou J *et al.* Simultaneous profiling of 3D genome structure and DNA methylation in single human cells. *Nat Methods* 2019;**16**:999–1006.

Lee S, Bakker CR, Vitzthum C *et al.* Pairs and Pairix: a file format and a tool for efficient storage and retrieval for Hi-C read pairs. *Bioinformatics* 2022;**38**:1729–31.

Li H. Aligning sequence reads, clone sequences and assembly contigs with BWA-MEM. *arXiv [q-bioGN]* 2013, DOI: 10.48550/ARXIV.1303.3997.

Liu H, Zhou J, Tian W *et al.* DNA methylation atlas of the mouse brain at single-cell resolution. *Nature* 2021;**598**:120–8.

Liu Z, Chen Y, Xia Q *et al.* Linking genome structures to functions by simultaneous single-cell Hi-C and RNA-seq. *Science* 2023;**380**:1070–6.

Martin M. Cutadapt removes adapter sequences from high-throughput sequencing reads. *EMBnet.journal* 2011;**17**:10–2.

Milo R, Jorgensen P, Moran U *et al.* BioNumbers--the database of key numbers in molecular and cell biology. *Nucleic Acids Res* 2010;**38**:D750–3.

Miura F, Enomoto Y, Dairiki R *et al.* Amplification-free whole-genome bisulfite sequencing by post-bisulfite adaptor tagging. *Nucleic Acids Res* 2012;**40**:e136.

Open2C, Abdennur N, Abraham S *et al.* Cooltools: Enabling high-resolution Hi-C analysis in Python. *PLoS Comput Biol* 2024a;**20**:e1012067.

Open2C, Abdennur N, Fudenberg G *et al.* Pairtools: From sequencing data to chromosome contacts. *PLoS Comput Biol* 2024b;**20**:e1012164.

Ramani V, Deng X, Qiu R *et al.* Massively multiplex single-cell Hi-C. *Nat Methods* 2017;**14**:263–6.

Tan L, Xing D, Chang C-H *et al.* Three-dimensional genome structures of single diploid human cells. *Science* 2018;**361**:924–8.

Tjong H, Li W, Kalhor R *et al.* Population-based 3D genome structure analysis reveals driving forces in spatial genome organization. *Proc Natl Acad Sci U S A* 2016;**113**:E1663–72.

Vasimuddin M, Misra S, Li H *et al.* Efficient architecture-aware acceleration of BWA-MEM for multicore systems. *2019 IEEE International Parallel and Distributed Processing Symposium (IPDPS)*. IEEE, 2019.

Wang X, Luan Y, Yue F. EagleC: A deep-learning framework for detecting a full range of structural variations from bulk and single-cell contact maps. *Sci Adv* 2022;**8**:eabn9215.

Wu H, Zhang J, Jian F *et al.* Simultaneous single-cell three-dimensional genome and gene expression profiling uncovers dynamic enhancer connectivity underlying olfactory receptor choice. *Nat Methods* 2024;**21**:974–82.

Wu P, Gao Y, Guo W *et al.* Using local alignment to enhance single-cell bisulfite sequencing data efficiency. *Bioinformatics* 2019;**35**:3273–8.

Yildirim A, Boninsegna L, Zhan Y *et al.* Uncovering the Principles of Genome Folding by 3D Chromatin Modeling. *Cold Spring Harb Perspect Biol* 2022;**14**, DOI: 10.1101/cshperspect.a039693.

Yildirim A, Hua N, Boninsegna L *et al.* Evaluating the role of the nuclear microenvironment in gene function by population-based modeling. *Nat Struct Mol Biol* 2023;**30**:1193–206.

Zhou T, Zhang R, Jia D *et al.* GAGE-seq concurrently profiles multiscale 3D genome organization and gene expression in single cells. *Nat Genet* 2024a;**56**:1701–11.

Zhou W, Johnson BK, Morrison J *et al.* BISCUIT: an efficient, standards-compliant tool suite for simultaneous genetic and epigenetic inference in bulk and single-cell studies. *Nucleic Acids Res* 2024b;**52**:e32.
